# Early EEG responses indicate liking for artworks presented in rapid visual streams

**DOI:** 10.64898/2026.09.24.754091

**Authors:** Sanjeev Nara, Vaishali Goyal, Gustavo Menegon, Philipp Flieger, Daniel Kaiser

## Abstract

When we appreciate visual art, we often take our time: We visit museums or galleries and deliberately engage with artworks and their meaning. Yet, our liking might at least partly be shaped by incidental encoding of the artworks’ visual features, a process initially independent from sustained engagement. Here, we show that liking for visual art can be predicted from rapid and automatic responses in the visual brain. We recorded EEG responses while participants viewed 999 artworks from the Vienna Art Picture System (VAPS), which features complex artworks that span different categories and art styles. Artworks were shown with a 300ms stimulus-onset-asynchrony (150ms on, 150 off), and participants responded to occasional target images. Using representational similarity analysis, we predicted the geometry of neural representations from the artworks’ similarity in liking (based on ratings from a different group of observers), while simultaneously controlling for similarities in other properties (category, art style, valence, arousal, and familiarity). Crucially, neural responses emerging within the first 150ms were robustly connected to liking, even though the task did not afford any liking judgments. Strikingly, this early neural correlate of liking was also observed in a second experiment, in which we presented the same 999 artworks with a 50ms stimulus-onset-asynchrony. Our results show that even under ultra-fast presentation conditions, automatic neural responses rapidly indicate liking. While fully appreciating an artwork may still need deliberate and prolonged engagement, the perceptual basis of whether we like or dislike an artwork seems to be laid in a split second.

## Introduction

When we go to museums or galleries, we take our time to evaluate an artwork. For instance, we may look at a painting for an extended amount of time, appreciate the meaning of what is depicted, think about the artist’s intentions, and relate the painting’s contents to our own experiences. All of these processes in turn influence whether we like the painting or not (*1-3*).

Yet, there may also be something fundamentally sensory about whether or not we like an artwork. Some compositions of visual features simply make more appealing artworks than others (*4, 5*). For instance, preferences for paintings depend on color compositions (*6*), the painting’s visual complexity (*7*), the balance between image elements (*8*), and on whether the composition follows favorable global configurations like the golden ratio (*9*).

Given the importance of visual features, neural responses in visual cortex should capture whether or not we ultimately like an artwork. Neuroimaging studies on the appreciation of artworks often highlight the involvement of higher-order valuation and reward systems in the frontal cortex or the default mode network (*10-13*). Yet, several studies also report that activity in multiple visual cortex regions, from low-level areas to category-selective modules in the ventral stream, encode the beauty of artworks (*11, 13, 14*). Such visual correlates of liking have been linked to favorable visual feature distributions that in turn reduce visual processing demands via sparser (*15*) or more fluent (*16, 17*) representations.

While these studies indicate that visual processes are involved in appreciating art, their participants typically provide explicit liking judgments. This renders it unclear whether the liking-related activations seen in visual cortex are indicative of differences in visual feature coding or whether they result from post-perceptual interactions across large-scale brain networks that emerge during the judgment of liking (*18*). To answer this question, time-resolved measures of brain activity are needed, ideally recorded in the absence of an explicit liking task.

Here, we recorded EEG responses while participants viewed 999 paintings from the Vienna Art Picture System (VAPS) database (*19*). In two experiments, we presented these artworks in rapid visual streams with stimulus-onset-asynchronies (SOAs) of 300ms (Experiment 1) and 50ms (Experiment 2), leaving little to no opportunities for cognitively engaging with the artworks’ content. In a representational similarity analysis (RSA) (*20*), we then related judgments of human liking for these artworks (judged by different observers in the VAPS database) to the EEG responses obtained under these rapid presentation regimes. In both experiments, EEG responses emerging within the first 150ms of processing reliably predicted liking, even when a range of control variables (e.g., the artworks’ style, category, valence, and arousal) were controlled for. These results demonstrate that liking is initially rooted in perceptual processing, even for complex visual artworks.

## Materials and Methods

### Participants

We conducted two EEG experiments. Experiment 1 was completed by 33 healthy adults (mean age 25 years, SD = 3.2; 15 female). Five additional participants were excluded due to not completing the experiment or due to excessive noise in the EEG recording. Experiment 2 was completed by a subset of 21 participants from Experiment 1 (mean age 26.1 years, SD = 3.5; 8 female). One additional participant was excluded due to excessive noise in the EEG recording. Participants received financial remuneration. All participants provided written informed consent. Procedures were approved by the ethics committee of the Justus Liebig University Giessen and were in accordance with the 6^th^ Declaration of Helsinki.

### Stimuli

The stimulus set consisted of 999 paintings, obtained from the VAPS database (*19*). This database includes a wide range of paintings that span five categories (landscapes, portraits, scenes, still lives, and towards abstraction) and 13 historical styles from the 15^th^ to the 21^st^ centuries (renaissance and mannerism, baroque and rococo, idealistic tendencies, realistic tendencies I (19^th^ century), realistic tendencies II (20^th^ to 21^st^ century), impressionistic tendencies, postimpressionistic tendencies, expressionistic tendencies, surrealistic tendencies, cubistic tendencies, constructivist tendencies, art informel tendencies, and abstract expressionist tendencies).

To account for the artworks’ varying aspect ratio, we generated 200 noise masks (square aspect ratio) using custom code based on mask images generated for continuous flash suppression experiments (https://martin-hebart.de/webpages/code/stimuli.html). Masks were generated in a way that approximately matched the color distribution and contrast of the artworks. During each trial of the experiment (see below), the artwork was then centrally overlayed on a randomly chosen mask image. This ensured that the stimulus on every trial of the EEG covered the same visual angle (Fig. 1A).

**Figure 1.**
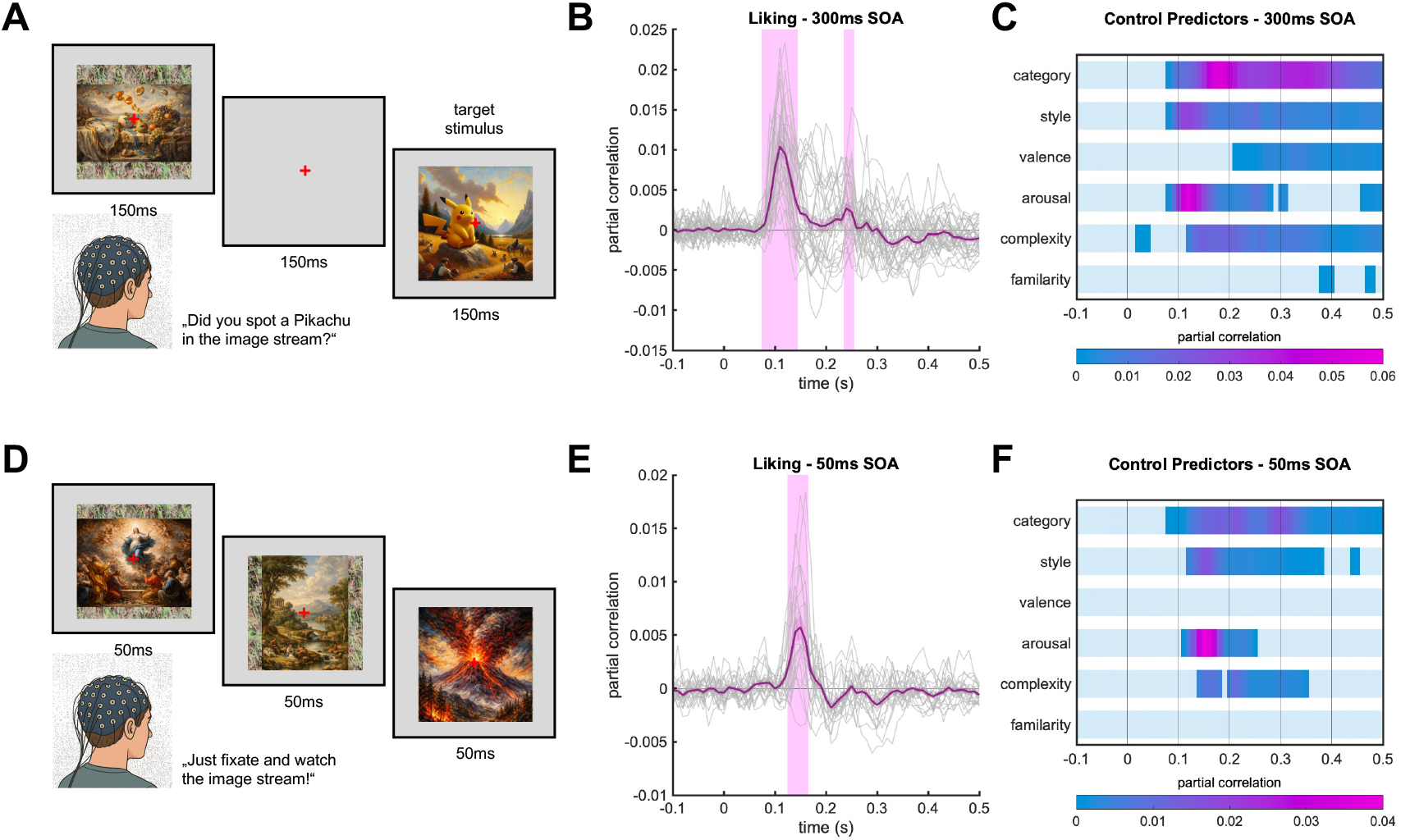
Rapid neural representations of liking. **A)** During Experiment 1, participants viewed the 999 VAPS artworks in streams of 50 images (300ms SOA) while we recorded their brain activity using EEG. After every stream, they were asked whether the stream did or did not contain a target images (an AI-generated artwork depicting Pikachu). Example paintings here and in (D) are generated with ChatGPT and were not in the actual stimulus set. **B)** Despite the rapid presentation regime and a task unrelated to aesthetics, liking predicted neural responses in the time periods from 90ms to 150ms and from 240ms to 250ms, even when the control predictors (see C) were accounted for. Gray lines represent individual participants, purple lines represent the average across participants, and pink shades indicate correlation significantly greater than 0. **C)** All control predictors also predicted neural responses, even if all the other predictors were partialed out. Only correlations significantly greater than 0 are highlighted. **D)** During Experiment 2, participants passively viewed the artworks in streams of 200 images (50ms SOA) while we again recorded their brain activity using EEG. **E)** Even under this ultra-rapid presentation regime, liking predicted neural responses from 130ms to 160ms. **F)** The artworks’ category and style, as well as arousal and complexity ratings also predicted neural responses. Together, these results show that fast-emerging neural responses during rapid visual stimulation are indicative of how much humans like an artwork.

The VAPS database also includes Likert-scale ratings for five properties of each artwork: liking (degree of preference), emotional valence (degree of emotional value), emotional arousal (level of excitement or stimulation), visual complexity (amount of the artworks’ details), and familiarity (how recognizable or well-known the artwork is), and. As our main focus was on liking, we focused our analyses on liking as the predictor of interest, while including the other ratings as well as the artworks’ categories and styles as control variables.

In addition, 20 computer-generated artworks featuring the Pokémon Pikachu were generated using the DeepAI image generator tool (https://deepai.org/machine-learning-model/text2img). Pikachu artworks were generated for each of the 13 art styles contained in the VAPS database. These images were used as target images in Experiment 1.

### Paradigm – Experiment 1

During Experiment 1, we presented the artworks in a series of rapid visual streams while recording EEG responses from participants. Participants were seated 60cm from the screen, with their head on a chinrest to reduce head movements. On each trial, a single artwork (5° × 5° visual angle) was presented for 150ms, followed by an inter-stimulus interval of 150ms. This amounts to a SOA of 300ms. A red fixation cross was shown in the center of the screen throughout to aid fixation. Stimulus presentation was controlled using the Psychtoolbox (*21*) in Matlab R2021a.

The experiment consisted of presentation streams of 50 trials. Each stream did or did not contain a target image (a Pikachu artwork; Fig. 1A). After every stream, participants were asked to indicate via mouse-click where a target was contained in the previous stream or not. Response options were presented on two half-circle segments that were randomly rotated for every response period. This task was designed to enforce processing of the artworks while being orthogonal to liking. Average task accuracy was 87.3% (SE=3.9%).

The experiment consisted of a total of 243 streams, during which each of the 999 artworks was repeated 12 times. The last stream was truncated after reaching the desired repetition count. The whole sequence was generated by shuffling the order of artworks 12 times and concatenating these shuffled orders, so that every artwork was shown once before the first repetition, and so on. The whole experiment lasted approximately 70 minutes.

### Paradigm – Experiment 2

In Experiment 2, we used the same paradigm with two changes. First, we further increased the presentation speed. Consecutive stimuli were now presented for 50ms, without any inter-stimulus interval, amounting to an SOA of only 50ms (*22*). Second, given this ultra-rapid presentation regime, we removed the task and let participants passively view streams of 200 artworks in a row (Fig. 1D). They could initiate the next stream at their own discretion by pressing the space bar.

The experiment consisted of a total of 60 presentation streams, during which each of the 999 artworks was again repeated 12 times. The whole experiment lasted approximately 10 minutes.

Experiment 2 was completed by a subset of 21 participants that also completed Experiment 1. These participants did both experiments in a single session, with 11 participants completing Experiment 1 first, and 10 participants completing Experiment 2 first.

### EEG recording and preprocessing

EEG data was recorded using a 64-channel Brain vision recorder with an Actichamp amplifier. Electrodes were placed according to the standard 10-10 system, using the Fz electrode as a reference. Data was recorded at 500 Hz. Preprocessing was performed offline using the Fieldtrip toolbox (*23*) in MATLAB R2024a. The continuous EEG data were epoched from -100ms to 1,000ms relative to stimulus onset, baseline-corrected for the pre-stimulus interval, and notch-filtered to remove 50Hz line noise. No low- or high-pass filters were applied. Noisy channels (max. 10 channels across participants) were rejected based on visual inspection, and eye blinks were removed using independent component analysis. Finally, the epoched data were resampled to 100Hz.

### Representational Similarity Analysis

We used RSA (*20*), implemented in CoSMoMVPA (*24*) for Matlab, to test how time-varying neural representations are influenced by the liking of the visual artworks. Specifically, we tested whether the representational dissimilarity among the 999 artworks in EEG response patterns across time (neural dissimilarity) is predicted by the dissimilarity in liking ratings for these artworks obtained from the VAPS database (liking dissimilarity). The following analyses were performed in an identical way for Experiments 1 and 2 and were conducted separately for each participant.

Neural dissimilarity was quantified by assessing correlations across electrode response patterns for each pair of artworks. For each time point along the EEG epochs (10ms resolution), we first performed principal component analysis (PCA) on the whole trials-by-electrodes array (retaining the components that explained 99% of the variance), in order to reduce the dimensionality of the feature space (*25*). We then averaged all trials on which the same artwork was shown, correlated feature vectors (i.e., response patterns across PCA components) across all possible pairs of artworks, and subtracted the resulting correlations from 1. Repeating this analysis for all time points across the epochs yielded a time course of neural representational dissimilarity matrices (RDMs), which indicate the artworks’ pairwise similarities in neural responses through time.

Liking dissimilarity was quantified by assessing differences in liking ratings from the VAPS database for each pair of artworks. Here, we computed the pairwise absolute differences in mean liking for each pair of artworks. This yielded a liking RDM, which indicates the artworks’ pairwise similarities in liking.

Akin to liking dissimilarity, we also quantified the artworks’ absolute rating differences in valence, arousal, complexity, and familiarity from the VAPS database, yielding four RDMs that indicated the artworks pairwise similarities in these properties. Further, we quantified whether the artworks came from the same or different categories and styles by constructing two RDMs that indicated whether an artwork features the same category/style (0) or a different category/style (1). These six RDMs were subsequently used as control variables, allowing us to uniquely attribute our results to liking.

To assess how liking is related to brain responses across time, we correlated (Spearman correlations) the neural RDMs and the liking RDMs at each time point across the epoch. To unequivocally attribute the results to a neural representation of liking, we used partial correlations, where correlations were assessed after partialing out the six control variables (category, style, valence, arousal, complexity, and familiarity). Further, we also assessed how the control variables were related to neural responses across time, by performing the same analysis with each of the control variables as the critical predictor while partialing out all other variables (including liking).

All analysis results were assessed in the time window between -100ms and 500ms. Given the rapid serial presentation regime with stimuli overlapping within each epoch, we did not expect any interesting results beyond 500ms post-stimulus.

### Statistical Analysis

Correlations between neural RDMs and predictor RDMs were compared to 0 using one-sided one-sample t-tests, performed separately at each time point between -100ms and 500ms. The resulting p-values were FDR-corrected to account for multiple comparisons across time. For each analysis, we report the peak timing as well as the test statistics (t-value and FDR-corrected p-value) at the peak.

### Data Availability

Data, code, and results files are publicly accessible on *zenodo* via this link: doi.org/ https://doi.org/10.5281/zenodo.21979554. Raw EEG data are accessible via this link: https://doi.org/10.5281/zenodo.22932789.

## Results

### Early EEG activity predicts liking in rapid visual streams

#### Experiment 1 – 300ms SOA

In Experiment 1, participants viewed 999 artworks in a rapid visual presentation regime with a SOA of 300ms, while engaged in a target detection task (Fig. 1A). Using RSA, we related neural responses across time to liking ratings obtained from the VAPS database.

Liking predicted neural responses from 80ms to 140ms and from 240ms to 250ms, with a peak at 110ms (t[32]=10.7, p_corr_<0.001). This indicated the fast-emerging neural responses, related to the perceptual processing of the artworks, are indicative of the aesthetic appeal of an artwork. Given that this neural signature of liking emerged under a rapid presentation regime and without a task related to liking, our results suggest that whether or not we like an artwork is partly explained by mandatory perceptual analysis in the visual system.

Critically, our analysis accounted for a range of control predictors, from the artworks’ category and style to ratings of valence, arousal, complexity and familiarity. This suggests that the observed responses are truly related to liking, and that liking can, for instance, be dissociated from the emotions that a painting elicits.

We also tested how the control predictors themselves predicted neural activity across time. All predictors were significantly related to neural responses. The artworks’ category predicted responses from 80ms to 500ms (peaking at 180ms, t[32]=11.7, p_corr_<0.001) and style predicted responses from 80ms to 500ms (peaking at 110ms, t[32]=15.3, p_corr_<0.001), suggesting that the most prominent style representations are reached before the most prominent category representations. Valence predicted responses from 210ms to 500ms (peaking at 330ms, t[32]=7.7, p_corr_<0.001) and arousal predicted responses from 80ms to 280ms, from 300ms to 310ms, and from 460ms to 500ms (peaking at 120ms, t[32]=14.3, p_corr_<0.001), indicating a fast and sustained representation of the emotional properties of an artwork that is distinct from and temporally more sustained than the rapid and transient representations that indicate liking. Further, the artworks’ arousal was more readily represented then their valence. Finally, complexity (from 120ms to 500ms; peaking at 150ms, t[33]=7.7, p_corr_<0.001; another significant time window from 20ms to 40ms is likely a false positive) and familiarity (from 380ms to 400ms and from 470ms to 480ms; peaking at 350ms, n.s.) also predicted neural responses to some extent.

These results also put the temporal profile of liking-related representations into context: Genuine representations of artwork liking emerged rapidly and were very transient in nature. Yet, this was not an artifact of the rapid serial presentation, as other properties were represented in a more sustained way even under this fast experimental regime.

#### Experiment 2 – 50ms SOA

In Experiment 2, we went one step further and presented the stimuli in ultra-rapid stimulus streams, with only 50ms SOA (Fig. 1D). We then conducted the same RSA as in Experiment 1 to test whether neural responses recorded during this ultra-rapid regime were still reliably related to liking.

Liking predicted neural responses from 130ms to 160ms, with a peak at 150ms (t[20]=5.0, p_corr_=0.001). This shows that even under an ultra-rapid presentation regime, early perceptual responses in visual cortex are indicative of whether or not an artwork is liked. The delayed peak relative to Experiment 1 may be indicative of a (forward-) masking effect caused by the rapid presentation regime.

As in Experiment 1, category (from 80ms to 500ms; peaking at 300ms, t[20]=4.5, p_corr_<0.001) and style (from 120ms to 380ms and 440ms to 450ms; peaking at 150ms, t[33]=9.9, p_corr_<0.001) predicted neural responses, too, and style-related responses again peaked earlier than category-related responses. Further, arousal (from 110ms to 250ms; peaking at 150ms, t[20]=9.8, p_corr_<0.001) and complexity (from 140ms to 180ms and from 200ms to 350ms; peaking at 210ms, t[20]=9.2, p_corr_<0.001) predicted neural responses, while valence and familiarity did not yield significant predictions in Experiment 2.

## Discussion

Here, we show that liking for visual artworks is partly reflected in early cortical responses linked to perceptual processing: Even under rapid serial presentation regimes, EEG responses emerging within 150ms after stimulus onset predicted human liking for a large set of paintings. This shows that despite the importance of non-visual factors such as deliberate engagement with, and emotional reflection on an artwork *(1-3, 26)*, visual processing is fundamentally related to preferences for visual art.

Neural representations related to liking emerged very fast, within the first 150ms of processing. This is distinctly faster than liking-related differences reported by previous M/EEG studies on visual art (*27-29*), which report the earliest effects after 300ms. There are two reasons for this discrepancy. First, using multivariate pattern analysis may provide higher power for detecting early effects, compared to the univariate analyses employed in the previous studies (*30*). Previous studies may thus have missed out on smaller effects during earlier time windows. Second, using a large stimulus set with hundreds of paintings provides a broad coverage of the visual feature space, which may be needed to cover the feature variations connected to liking. The smaller stimulus sets used in previous studies may simply not tap into the relevant feature dimensions to the required extent. Indeed, the neural correlates of liking for large sets of real-world objects also emerge in a similarly rapid fashion (*31*).

Neural representations related to liking were also surprisingly transient. Unlike other properties like an artwork’s category and style or its emotional associations, liking was only represented briefly around 100ms to 150ms after onset and later responses did not reliably reflect liking. The early perceptual processes that shape representations during this transient time period may indicate feature-based precursors of liking. These early precursors may form relatively stubborn representations that are extracted incidentally (*31, 32*), even during passive viewing, and may not be overridden by prolonged viewing and engagement when asked for liking judgments. This may not hold true for later perceptual analysis: Once more time is available for evaluating the image, more complex visual feature attributes (reflected in later neural responses) may be flexibly re-weighted during more sustained engagement with a stimulus when liking is explicitly judged. An alternative is that the continuous masking in our rapid visual presentation regime interrupts recurrent interactions in visual cortex (*33, 34*), which may be needed to generate the feature representations indicative of liking.

We could still reliably predict liking when neural responses were recorded under a 50ms SOA regime, where stimuli are just barely visible to participants due to excessive forward- and backward-masking. While this is consistent with visual representations of other properties like contrast, color, and object category emerging reliably under such fast presentation regimes (*22, 35*), it further stresses that such fast presentation regimes can also be used to distill the neural correlates of higher-level perceptual decisions like whether or not we like an artwork. It is worth noting that the whole of Experiment 2 took only 10minutes. Future studies could therefore use similar rapid serial presentation paradigm to provide a quick and coarse neural benchmarking of visual preferences in a given set of candidate images.

In our study, we could predict liking for artworks from neural responses that emerge during a liking-unrelated task (Experiment 1) or during passive viewing (Experiment 2). This is consistent with findings for natural photographs, where artificial neural networks that are not trained to evaluate the beauty of an image still provide powerful predictions of beauty (*15, 16, 36, 37*). Also for artworks, artificial neural networks trained on image categorization can provide good models for human liking (*17, 37*). This suggests that the incidental and automatic representation of visual features already provides rich information for downstream systems more directly involved in liking judgments.

Our approach of predicting liking judgments provided by one group of participants from neural responses recorded for another group of participants inherently relies on participants agreeing on what is beautiful. Yet, the aesthetic appeal of paintings can vary drastically across individuals (*38*). Our study therefore provides insights into how visual brain responses determine liking at the group level and may miss out on the idiosyncrasies in both liking and its neural correlates. As even early cortical responses related to beauty have been shown to vary across participants (*39, 40*), it would be interesting to see whether the rapid and automatic neural representations uncovered here can predict idiosyncrasies in beauty judgments.

Finally, it is only fair to stress that appreciating art is far more than an evaluation of simple visual features. When we take our time to fully appreciate an artwork, these early visual representations are likely read out, remapped, and interpreted across multiple brain systems (*11, 18, 41, 42*), finally resulting in an aesthetic evaluation. Across this entire processing cascade, a plethora of cognitive processes ultimately shape our enjoyment of a piece of art. Yet, this whole process starts with favorable feature combinations that are represented rapidly and automatically during visual processing.

## Acknowledgements

This project was supported by the German Research Foundation (DFG), grant KA4683/6-1 (project number 536053998) and under Germany’s Excellence Strategy (EXC 3066/1 “The Adaptive Mind”, Project No. 533717223).

## Author contributions

Conceptualization: SN and DK

Data curation: SN, VG, and GM

Formal analysis: SN, VG, GM, PF, and DK

Funding acquisition: DK

Investigation: SN, VG, and GM

Methodology: SN and DK

Project administration: SN, PF, and DK

Resources: SN and DK

Software: SN, VG, GM, PF, and DK

Supervision: SN, PF, and DK

Validation: SN, VG, GM, and DK

Visualization: DK

Writing—original draft: DK

Writing—review and editing: SN, VG, GM, PF, and DK

